# The zebrafish as a new model for studying chaperone-mediated autophagy unveils its role in spermatogenesis

**DOI:** 10.1101/2024.06.10.597508

**Authors:** Maxime Goguet, Emilio J Vélez, Simon Schnebert, Karine Dias, Vincent Véron, Alexandra Depincé, Florian Beaumatin, Amaury Herpin, Iban Seiliez

## Abstract

Chaperone-Mediated Autophagy (CMA) is a major pathway of lysosomal proteolysis involved in numerous cellular processes, and whose dysfunction is associated to several pathologies. Initially studied in mammals and birds, recent findings have identified CMA in fish, reshaping our understanding of its evolution across metazoans. Given the exciting perspectives this finding offered, we have now developed the required tools to investigate and functionally asses that CMA function in a powerful fish genetic model: the zebrafish (*Danio rerio*). After adapting and validating a fluorescent reporter (KFERQ-Dendra2; previously used to track CMA in mammalian cells) in zebrafish primary embryonic cells, we first demonstrated CMA functionality in this fish species. Then, we developed a transgenic zebrafish line expressing the KFERQ-Dendra2 CMA reporter, enabling the real-time tracking of CMA activity *in vivo*. This model revealed heterogeneous CMA responses within tissues, highlighting the zebrafish as a valuable model for investigating tissue-specific and cell-scale variations in CMA. Moreover, a novel role for CMA has been uncovered, acting as a gatekeeper of sperm cell proteostasis, thereby playing a crucial role in the production of active and high-quality spermatozoa. Overall, these findings emphasize the zebrafish as a pivotal model for advancing our comprehension of the fundamental mechanisms underlying CMA.

## INTRODUCTION

Chaperone-Mediated-Autophagy (CMA) is a major pathway of lysosomal proteolysis essential for the control of cellular homeostasis and metabolism^1^. Due to its ability to target damaged or non-functional proteins for degradation, CMA has been shown to be critical for intracellular protein quality control^2^. Another core feature of CMA is the wide diversity of proteins it selectively targets, which connects this function to the control of numerous intracellular processes, including (but not limited to) glucose and lipid metabolism^3^, cell cycle^4^, transcriptional programs^5^, immune response^6^ and pluripotency of stem cells.^7^ Unsurprisingly, CMA dysfunctions have been associated to the occurrence of several human pathologies, including neurodegenerative diseases, cancers and immune disorders^8^. Current research efforts are now underway to better clarify the underlying canonical mechanisms.

In details, during CMA, cytosolic proteins containing a pentapeptide sharing biochemical similarity to the KFERQ (lysine-phenylalanine-glutamate-arginine-glutamine) sequence are first recognized by the heat-shock protein family A [Hsp70] member 8 (HSPA8/HSC70) and co-chaperones^9,10^. The substrate-chaperone complex then docks at the lysosomal membrane through specific binding to the cytosolic tail of the lysosomal associated membrane protein 2A (LAMP2A), the limiting and essential protein for CMA^11^. LAMP2A then organizes into a multimeric complex that allows the substrate to translocate across the lysosomal membrane^12,13^, where it is degraded by acid hydrolases.

Initial studies on CMA mainly focused on *in vitro* assessments of the different steps of the process on isolated lysosomes^14–17^. Although these experiments resulted in major advances for our general understanding regarding to the detailed processes involved in CMA as well as their regulations at the whole tissue level, they nevertheless failed to provide comprehensive insights at the cellular level and/or to discriminate cell-type differences in CMA. A significant step forward in the field was then reached with the development and validation of a fluorescent CMA reporter system, which allowed monitoring CMA activity in living cells^18^. Such a CMA reporter could be used in a broad variety of cells, and it became clear that levels of basal and inducible CMA activity are indeed cell-type dependent^18^. However, the use of this reporter was limited to *in vitro* models, and studies on animal tissues, which still relied on the measurement of CMA activity on isolated lysosomes, remained unanswered regarding to cell type-dependent variations in CMA within complex tissues. A major step forward in CMA research came with the recent development and validation of a transgenic mouse model ubiquitously expressing a fluorescent CMA reporter (KFERQ-Dendra2) that allowed monitoring and quantifying this process *in vivo* at a cellular resolution^19^. Although generated only recently, this model already contributed to a significant more in-depth understanding of the properties and regulations of this form of autophagy^19–22^.

Alongside these technical advances, the recent discovery of the existence of CMA in fish^23,24^ opened up new and exciting avenues in the field through the possibility of now using complementary and powerful genetic models, such as zebrafish or medaka, for studying this fundamental function also in an evolutionary perspective. Particularly, it allows benefiting from the numerous advantages offered by these models (high number of offspring, transparency of the young stages allowing live fluorescent imaging during organogenesis, fully sequenced genome, ease of forward/reverse genetics…) to expand our basic knowledge of CMA at multiple organismal levels, including that of single cell, whole tissues and organism, as already demonstrated in many fields of biology^25^.

In this context, the aim of the present study was to determine to what extend these expectations are grounded, and thus to validate the zebrafish as a relevant, powerful and central organism complementary to murine models for studying CMA. The fluorescent CMA reporter system - previously developed to track CMA in mammalian cells^18,19^ - was first tested in zebrafish primary embryonic cells. Our results show that upon starvation and/or mild oxidative stress, that CMA reporter re-localizes from a diffuse cytoplasmic distribution to lysosomes in a KFERQ- and Lamp2a-dependent manner, hence providing unequivocal evidence for the existence of a functional CMA process in this fish species, as previously demonstrated in medaka fish^23^ and more recently in rainbow trout^26^. Further on, we established a KFERQ-Dendra2 transgenic reporter zebrafish line and demonstrated that these fish allow – within tissues – the live-tracking, discrimination and quantification of cell-type variations between basal and inducible CMA activity. All in one, these technical improvements notably allowed monitoring the so far un-reported spatial distribution of a CMA activity in the testes, and revealed an unforeseen role for this function as a gatekeeper of sperm cell proteostasis and quality, ultimately contributing to the production of active spermatozoa. Collectively, these results validate the KFERQ-Dendra2 transgenic zebrafish as a reliable model to improve our understanding of the pathophysiology and mechanisms underlying the CMA (dys)functions.

## RESULTS

### Zebrafish cells recapitulate a whole CMA process from KFERQ-motif recognition to lysosome delivery and degradation in a Lamp2a-dependant manner

To substantiate and definitively apprehend CMA or a CMA-like process in fish we first adapted and validated - in zebrafish - the use of a photo-switchable (PS) fluorescent CMA reporter (KFERQ-PS-Dendra2), which has proven to be a reliable reporter for tracking and measuring CMA activity in mammals^18,19^. This reporter consists of the N-terminal 21 amino acids of bovine RNase A, including its KFERQ-CMA targeting motif (highly conserved among vertebrates), fused to the PS fluorescent protein Dendra2. Activation of CMA activity (for instance after prolonged starvation or under oxidative stress) induces re-localization of the fluorescent reporter from a cytoplasmic diffuse distribution to clearly identifiable and easily quantifiable *puncta* that co-localize within lysosomes^18^. After micro-injection of KFERQ-PS-Dendra2-coding mRNAs in just fertilized zebrafish zygotes, cells from 50% epiboly staged embryos were isolated and photo-activated with a 405 nm light emitting diode for 10 min (Figure 1A). Cells were then maintained for 16h with (Serum+) or without serum (Serum-), in presence or absence of hydrogen peroxide (H_2_O_2_). In control condition (Serum+), the reporter appeared diffusely localized throughout the cytoplasm with no or very few *puncta* detected (Figure 1B; quantification in Figure 1C). In contrast, after starvation (Serum-) or upon H_2_0_2_ treatment or both (Serum+; H_2_O_2_ and Serum-; H_2_O_2_), cells displayed a significantly higher number of KFERQ–Dendra2 *puncta* (Figure 1B; quantification in Figure 1C), co-localizing together with the lysosomal marker Lamp1 (Figure 1D; quantification in Figure 1E).

**Figure 1.**
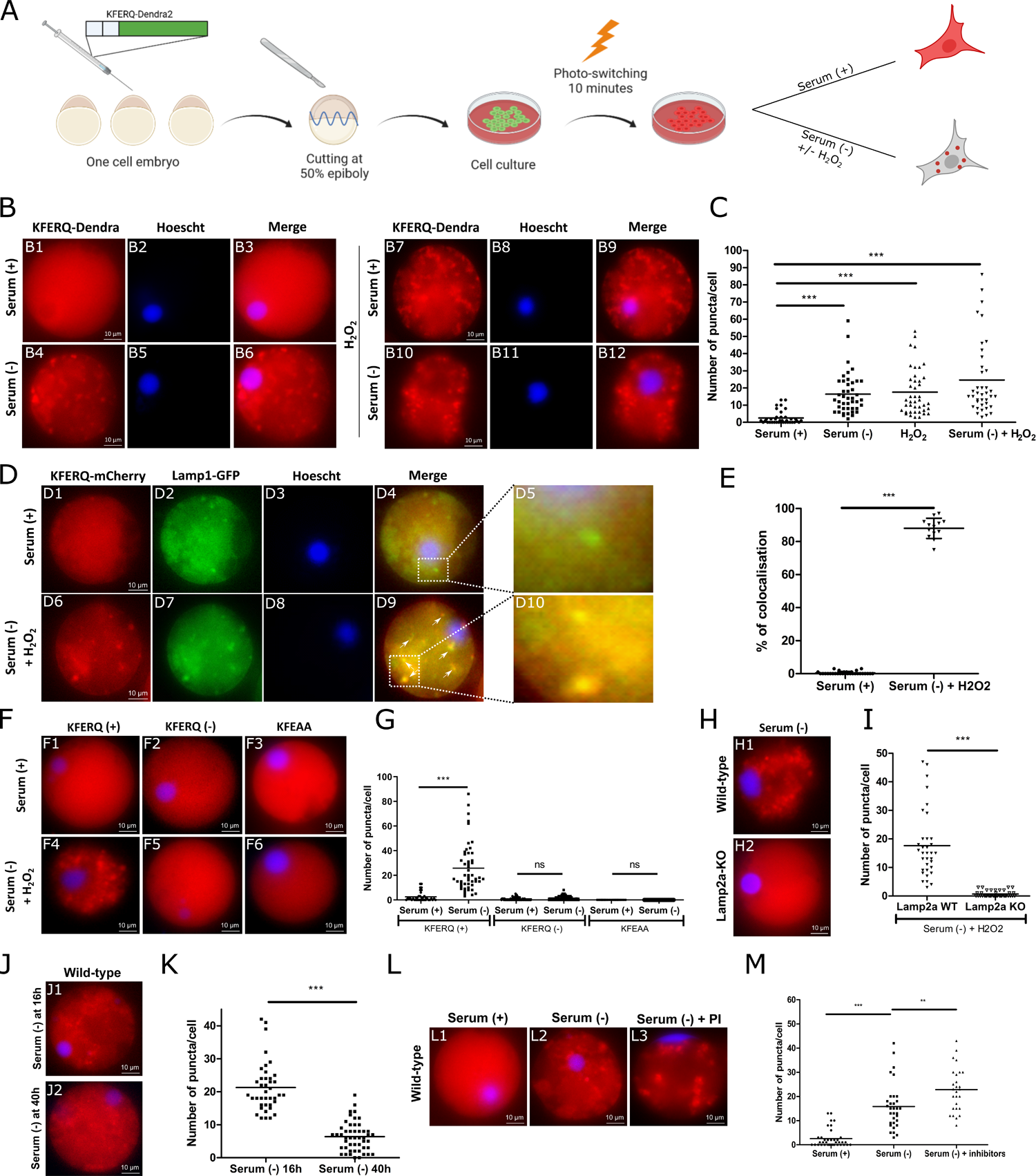
Zebrafish primary embryonic cells display a functional CMA process. (A) Experimental Setup. Microinjection of KFERQ-Dendra2-coding mRNAs in just fertilized zebrafish zygotes. Cells from 50% epiboly embryos were isolated and photoactivated with a 405lJnm LED for 10 minutes. Cells were then subjected to various conditions: untreated (Serum+), mild oxidative stress (H_2_O_2_, 25µM), and/or serum deprivation (Serum-) for 16 h. CMA activity was quantified by counting KFERQ-Dendra2 puncta. (B) Representative images of zebrafish primary embryonic cells visualized by super-resolution fluorescence microscopy (scale bar, 10 µm). Nuclei were stained with Hoescht 33342 (blue). Under serum-rich conditions (Serum+), the reporter diffusely localized in the cytoplasm with few *puncta* (B1 to B3). In serum-deprived and/or H_2_O_2_-treated conditions, *puncta* patterns were observed. (B4 to B12). (C) Quantification of KFERQ-Dendra2 reporter *puncta* per cell (n=50 cells/condition). Averages (horizontal lines) obtained with cells cultured without serum (Serum-) and/or with H_2_O_2_ were compared with those of control cells (Serum+) using a Student t-test (***, p<0.001). (D) Representative images of zebrafish primary embryonic cells microinjected with KFERQ-PA-mCherry1 (red) and LAMP1-GFP (green). Under Serum+ condition, the CMA reporter is diffusely localized throughout the cytoplasm (D1) and no co-localization with LAMP1-GFP is observed (D4 and D5). However, upon serum deprivation and H_2_O_2_ treatment, most of the KFERQ-PA-mCherry1 *puncta* (D6) co-localized with the green lysosomal marker LAMP1-GFP (D7), as indicated by the arrowheads (D9) and amplified in the insets (D10). (E) Quantification of percentage of co-localization of CMA reporter *puncta* with LAMP1-GFP per cell under Serum+ and Serum-/H_2_O_2_ conditions (n=50 cells/condition). A Student t-test was performed to compare both conditions (***, p<0.001). (F) Representative images of zebrafish primary embryonic cells micro-injected with mRNAs coding for either the CMA reporter KFERQ-Dendra2 (KFERQ+) (F1 and F4), the PA– Dendra2 construct lacking the KFERQ motif (KFERQ-) (F2 and F5), or a mutant reporter with a non-functional KFEAA-sequence (KFEAA) (F3 and F6). After photoactivation, cells were cultivated in Serum+ (F1 to F3) or Serum-/H_2_O_2_ (F4 to F6) conditions. Nuclei were stained with Hoescht 33342 (blue). (G) Quantification of the number of *puncta* obtained with each construction and under each condition described in F. All values correspond to individual images (n=50 cells/condition). The lines represent the average for each condition. For each construction, the Serum+ condition was compared to the Serum-/H_2_O_2_ condition with a t-test (***, p<0.001). (H) Representative images of primary embryonic cells from WT (H1) or Lamp2a-KO (H2) zebrafish micro-injected with mRNAs coding for the KFERQ-Dendra2 reporter. Cells were maintained in the Serum-/H_2_O_2_ condition for 16 h. Nuclei were stained with Hoescht 33342. (I) Quantification of KFERQ-Dendra2 reporter red puncta per cell. All values correspond to individual images (n=50 cells/condition). The lines represent the average for each condition. The two conditions were compared with a Student t-test (p<0.001***). (J) Representative images of zebrafish primary embryonic cells micro-injected with mRNAs coding for the KFERQ-Dendra2 reporter and maintained in the absence of serum (Serum-) for 16 h (J1) and 40h (J2). (K) Quantification of KFERQ-Dendra2 reporter red puncta per cell. All values correspond to individual images (n=50 cells/condition). The lines represent the average for each condition. The two conditions were compared with a t-test (p<0.001***). (L) Representative images of zebrafish primary embryonic cells micro-injected with mRNAs coding for the KFERQ-Dendra2 reporter and maintained in a serum-rich medium (Serum+), serum-free medium (Serum-) or serum-free medium containing a lysosomal protease inhibitor (Serum- / PI) for 16h. (M) Quantification of KFERQ-Dendra2 reporter red puncta per cell. All values correspond to individual images (n=50 cells/condition). The lines represent the average for each condition. The different conditions were compared with a t-test (p<0.001***).

In order to determine whether the formation of these observed *puncta* required the KFERQ-targeting motif, zebrafish cells were then micro-injected with mRNAs encoding either a PA–Dendra2 construct lacking the KFERQ motif or a mutant reporter with a non-functional KFEAA-sequence replacing the KFERQ motif^27^. Compared with the KFERQ-Dendra2 reporter (KFERQ (+)), both the Dendra2 alone (KFERQ (-)) and the mutant KFEAA-Dendra2 (KFEAA) fluorescence remained mainly diffusely localized throughout the cytoplasm and no *puncta* could be observed after prolonged starvation (Figure 1F; quantification in Figure 1G). This demonstrated that the formation of *puncta* is contingent to the presence of the KFERQ-motif.

In mammals, KFERQ-domain bearing proteins are additionally subjected to parallel degradation in late endosomes/multivesicular bodies through a pathway referred as <u>e</u>ndosomal <u>MI</u>croautophagy (eMI)^28^. However, unlike CMA, this process does not require the LAMP2A protein, but relies on the “Endosomal Sorting Complex Required for Transport” (ESCRT) machinery^28^. Hence, to determine the mechanisms accounting for the *puncta* formation observed with our KFERQ-Dendra2 reporter, we therefore monitored the activity of the reporter in fish cells specifically depleted of Lamp2a (Lamp2a-KO, unable to perform CMA; generated using the genome-editing CRISPR-Cas9 tool (Figure S1A)). In details, the produced Lamp2a-mutant fish displays a deletion of a 737-bp region in exon 6 of the *lamp2* gene that encodes for the specific cytosolic and transmembrane domains of the Lamp2a protein (Figures S1B and S1C). Levels of *lamp2a* mRNAs were undetectable in these fish, whereas those of the two others *lamp2* splice-variants transcripts (*lamp2b* and *lamp2c*) were comparable with their wild-type counterparts (Figure S1D). After 16 h starvation (serum-), the KFERQ-reporter appeared diffusely localized throughout the cytoplasm of the Lamp2a-KO cells and no *puncta* was detectable (Figure 1H; quantification in Figure 1I).

Starvation and H_2_O_2_ activate macroautophagy^29–31^. To determine whether the observed *puncta* could be attributed to a possible macroautophagy activity, we next used antisense morpholino oligonucleotides (Mo) to inhibit translation of zebrafish mRNAs encoding the E1-like enzyme Atg7, a core macroautophagy-related factor involved in the conjugation of Atg5 to Atg12^32^. Lowering *atg7* expression after Mo injection led to the loss of Atg12–Atg5 complex formation (Figures S2A and S2B), but did not affect the induction of *puncta* by 16 h starvation (Figure S2C; quantification in Figure S2D).

The entire CMA process does not only imply the recruitment of the KFERQ domain containing substrate to the lysosomal membrane but also its translocation and subsequent degradation within the lumen. To assess the correct lysosomal internalization and degradation of the CMA reporter upon starvation, we next monitored the dynamics of changes in the number of fluorescent *puncta*, which is expected to decrease as the protein is internalized and degraded in this compartment. The strong reduction in the number of puncta/cell observed after long-term serum deprivation (40h) compared to cells starved for a shorter time (16h) supports that the fluorescent reporter was indeed internalized and degraded over time (Figures 1J1 and 1J2, quantification in Figure 1K). Into that direction, inhibition of lysosomal proteases through leupeptin+NH_4_Cl treatment induced a significant increase in the number of *puncta* upon serum depletion (Figures 1L1-1L3, quantification in Figure 1M).

Together, these results support the existence, in zebrafish, of a pathway that, upon fasting and/or oxidative stress, specifically targets KFERQ-proteins in a Lamp2a-dependent manner for lysosomal degradation, similarly to the previously defined CMA process in mammals and more recently in medaka^23^ and rainbow trout^26^.

### The zebrafish KFERQ-Dendra2 line allows for CMA monitoring and quantification *in vivo* at a cellular resolution

The characterization of a CMA activity in zebrafish now opens up the possibility of using this species - a powerful model organism for functional/developmental studies - to improve our fundamental understanding of this function across vertebrates. To this end, we generated a transgenic zebrafish line ubiquitously expressing the fluorescent CMA reporter KFERQ-Dendra2 described above and which, in mammals, has been proven to be a reliable reporter for monitoring and measuring CMA activity *in vivo* as well^19^. In details, we used a vector suitable for *in vivo* expression using the 3167 plasmid backbone (see materials and methods for details) and inserting the oligonucleotide coding for the N-terminal sequence of the bovine ribonuclease A containing the KFERQ-CMA targeting motif in frame with the sequence of the photo-convertible fluorescent protein Dendra2 under the ubiquitin promoter that provides ubiquitous expression (Figure 2A). In absence of photoactivation, the KFERQ-Dendra2 fish ubiquitously express a green fluorescence (emission collected at 507 nm after 488 nm excitation) while no red fluorescence is observable (emission collected at 576 nm after 561 nm excitation) (Figures 2B1 and 2B2). In contrast, following photoactivation at 405 nm for 10 minutes, the KFERQ-Dendra2 fish emitted also a uniformly distributed red fluorescence in all tissues examined (Figures 2B3-2B8).

**Figure 2.**
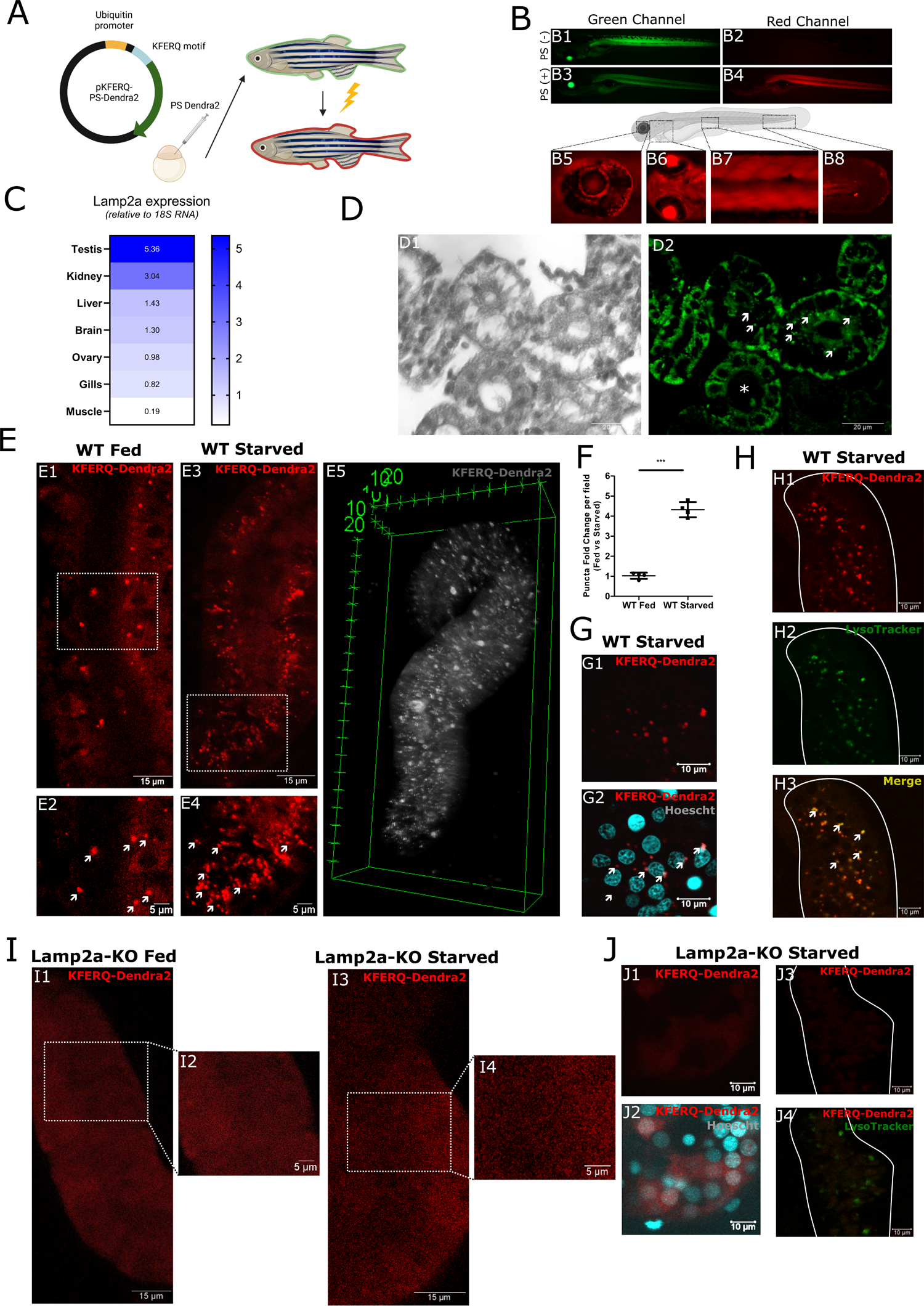
Monitoring CMA activity at cellular resolution using the zebrafish KFERQ-Dendra2 line. (A) Left: Map of the vector used for generation of the KFERQ-PS-Dendra2 transgenic zebrafish line. Right: Experimental Setup. (B) Zebrafish KFERQ-Dendra2 line were exposed (Photoswitch+; PS+) or not (Photoswitch-; PS-) to a 405nm light for 10 min, and photoswitching was monitored by imaging fish in the green (B1 and B3) and red (B2 and B4) channels. Following exposure to 405 nm light, KFERQ-Dendra2 fish emit red fluorescence in all tissues examined (B5-B8). B5: eye in lateral view; B6: head in ventral view; B7: trunk in lateral view; B8: tail in lateral view. (C) RT-qPCR for lamp2a in the indicated tissues relative to 18S RNA in starved conditions, n=3 fish. (D) Brightfield (D1) and green fluorescence (D2) imaging of kidney sections from KFERQ-Dendra2 zebrafish starved for 48 h. Green Dendra positive puncta (arrows) are shown. Asterisk shows the collecting duct. (E) Representative images of kidney tubules from fed (E1) or 48 h starved (E3) KFERQ-Dendra2 zebrafish. E2 and E4 show boxed areas at higher magnification. E5 shows a 3D representation of a tubule from KFERQ-Dendra2 zebrafish starved for 48 hours. (F) Dendra-positive puncta quantification from 4 kidneys per line from 4 independent experiments. (G) Red KFERQ-Dendra2 fluorescence (G1) and Hoechst nuclear staining (G2) in kidney tubules from 48 h starved KFERQ-Dendra2 zebrafish. (H) Red fluorescence (H1) and Lysotracker staining (H2) of kidneys from 48 h starved KFERQ-Dendra2 zebrafish reveal clear co-localization of the two signals (arrows in H3), confirming the lysosomal localization of the reporter. (I) KFERQ-Dendra2 red fluorescence in kidney from fed (I1) or starved for 48 h (I3) KFERQ-Dendra2-Lamp2a-KO zebrafish. I2 and I4 show boxed areas at higher magnification. (J) Red KFERQ-Dendra2 fluorescence (J1) and Hoechst nuclear staining (J2) in kidney tubules from 48 h starved KFERQ-Dendra2-Lamp2a-KO zebrafish. (J3) Red fluorescence and (J4) Lysotracker staining of kidneys from 48 h starved KFERQ-Dendra2-Lamp2a-KO zebrafish reveal no co-localization of the two signals.

As a proof of concept for being able to visualize fine variations in CMA activity at the cellular level within different tissues using the zebrafish KFERQ-Dendra2 reporter line, we first sampled kidneys – an organ known to exhibit a high basal and induced CMA activity in mammals^19^. Interestingly, a high *lamp2a* mRNA expression is also observed in zebrafish (Figure 2C). Analysis of the kidney from the KFERQ-Dendra2 zebrafish, revealed the presence of discrete CMA reporter-positive *puncta* in tubules (Figures 2D1 and 2D2). In details, we found that, in fed control fish, these *puncta* were sparsely detected (Figures 2E1 and 2E2). In contrast, kidneys sampled from starved fish displayed a significantly higher number of KFERQ–Dendra2 *puncta* in tubules (Figures 2E3-2E5 and video S1; quantification in Figure 2F), mainly located at the perinuclear region (Figures 2G1 and 2G2), known to be abundant in lysosomes^33,34^. In this regard, we found that the fluorescent CMA *puncta* co-localized together with the LysoTracker (Figures 2H1-2H3), supporting a lysosomal-restricted localization of the reporter. We also confirmed that the expression of KFERQ-Dendra2 mRNA remained unchanged upon starvation, indicating that the changes observed in the number of *puncta* between fed and starved fish were not a consequence of any possible changes in the expression of the CMA reporter (Figure S3A-C).

In order to provide further validation of the selectivity of the reporter zebrafish model for CMA, we then crossed KFERQ-Dendra2 zebrafish with our Lamp2a-KO line. Kidney from KFERQ-Dendra-Lamp2a-KO fish did not display any visible fluorescent *puncta* neither in fed nor fasted conditions (Figures 2I1 – 2J2 and Video S2), while the total number of the lysosomes in these cells was not affected (Figures J3 and J4). Finally, we could also confirm similar expression levels of the KFERQ-Dendra2 mRNA between the WT and Lamp2a-KO CMA reporter fish, indicating that the absence of *puncta* in the latter is not due to a lack of CMA reporter transgene expression (Figure S3).

Together, these data demonstrate that the KFERQ-Dendra2 protein expressed in our CMA reporter zebrafish line relocates upon starvation from a cytoplasmic diffuse distribution to *puncta* that co-localize with lysosomes in a Lamp2A-dependent manner as a consequence of an active CMA process. As such, this zebrafish KFERQ-Dendra2 line allows monitoring cell type variations in basal and inducible CMA within the same tissue.

### A new and key role for CMA during spermatogenesis

Although increased levels of LAMP2A and HSC70 have been reported in the testes of the *Stra8-cre;Foxj2^tg/tg^* sterile mice model^35^, the possible involvement of CMA in spermatogenesis has yet not been investigated in details. Analysis of the testes of KFERQ-PS-Dendra zebrafish showed low basal CMA activity in this tissue (Figures 3A1-3A3). However, after starvation, a 3-fold increase in the number of fluorescent *puncta* was recorded (Figures 3A4 – 3A6; quantification in Figure 3A7). As evidenced by the Tg(*gsdf*:eGFP);KFERQ-PA-mCherry zebrafish line, a strong regionalization of CMA activity within the testis was also observed, with *puncta* occurring exclusively within specific cells known as Sertoli cells (eGFP positive) that surround the germ cells (Figures 3B1-3B3). Interestingly, a recent evolutionary-driven study reporting single-nucleus transcriptome data from testes of eleven species covering the three major mammalian lineages (eutherians, marsupials, egg-laying monotremes) and birds, and including seven key primates, showed that the *lamp2* gene is predominantly expressed in Sertoli cells in all species analyzed (except in mouse, which show a different expression profile). More than reporting an evolutionary conservation of the expression profile of this gene across vertebrates, these results indeed suggest or question about a role for CMA in testes physiology, that could be accomplished by the Sertoli cells (Figure S4 reproduced with permission from Murat et al.^36^).

**Figure 3.**
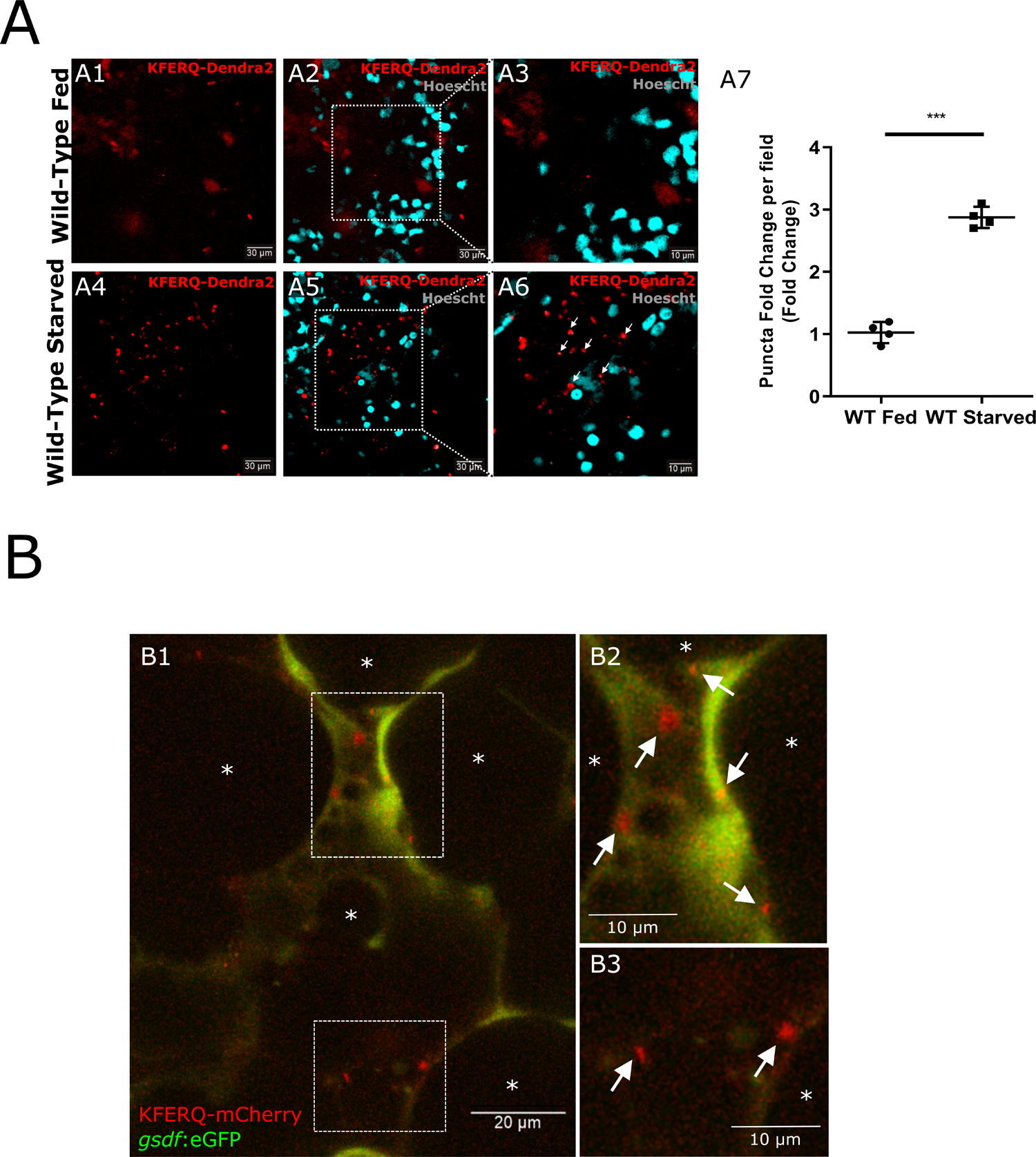
CMA activity in testes. (A) Red KFERQ-PS-Dendra2 fluorescence (A1 and A4) and Hoechst nuclear staining (A2 and A5) in testes from fed (A1 and A2) and 48 h starved (A4 and A5) KFERQ-PS-Dendra2 zebrafish. A3 and A6 show boxed areas at higher magnification. A7, Dendra-positive *puncta* quantification from 4 testes per line from 4 independent experiments. (B) Red KFERQ-PA-mCherry + green *gsdf*:eGFP fluorescence in testis from the Tg(*gsdf*:eGFP);KFERQ-PA-mCherry zebrafish transgenic line. The proximal *gsdf* (*gonadal soma-derived factor*) gene promoter drives eGFP expression specifically in the Sertoli cells. Red KFERQ-PA-mCherry positive *puncta* (arrows) and spermatogenic cysts containing germ cells (*) are shown. B2 and B3 show boxed areas at higher magnification.

Given the major role of Sertoli cells in orchestrating and supporting the development of germ cells into mature spermatozoa in the adult male gonad^37^, we sought to determine the consequences of Lamp2a invalidation on testicular development and function. H&E staining of testes sections from WT and Lamp2a-KO fish first revealed that the latter exhibit significantly higher sperm counts than their WT counterparts (Figures 4A1-4A8; quantification in Figure 4B). We then performed a comparative transcriptomics of testes of both lines, and found that mRNA expression levels of different factors involved in dynein-complex assembly (DNAAF3, DNAI3/WDR63) and actin filament remodeling (Eps8l1a, lima1a) differed significantly between Control and Lamp2a-KO fish (Figure 4C and Table S1). These first results, prompted us to assess sperm mobility characteristics of our Lamp2a-KO fish, as defects in the assembly of axonemal dynein arms as well as in actin filament remodeling have been shown to cause male infertility due to asthenozoospermia^38,39^. To that end, spermatozoids from either WT or Lamp2a-deficient fish were subjected to the <u>c</u>omputer-<u>a</u>ssisted <u>s</u>perm <u>a</u>nalysis (CASA) protocol system in order to provide objective and precise information on the kinetic of sperm cells. Interestingly, a drastic reduction in the percentage of motile spermatozoids from Lamp2a-KO fish compared to those from WT was recorded (Figures 4D1-4D3 and Videos S3-S4). Additionally, several other parameters including the average path velocity (VAP), the curvilinear velocity (VCL) as well as the straight velocity (VSL) consistently gave lower values for Lamp2a-KO sperm when compared to its WT counterpart (Figure 4E1 - 4E4). Taken together, these findings reveal a previously unknown role for CMA in producing active sperm.

**Figure 4.**
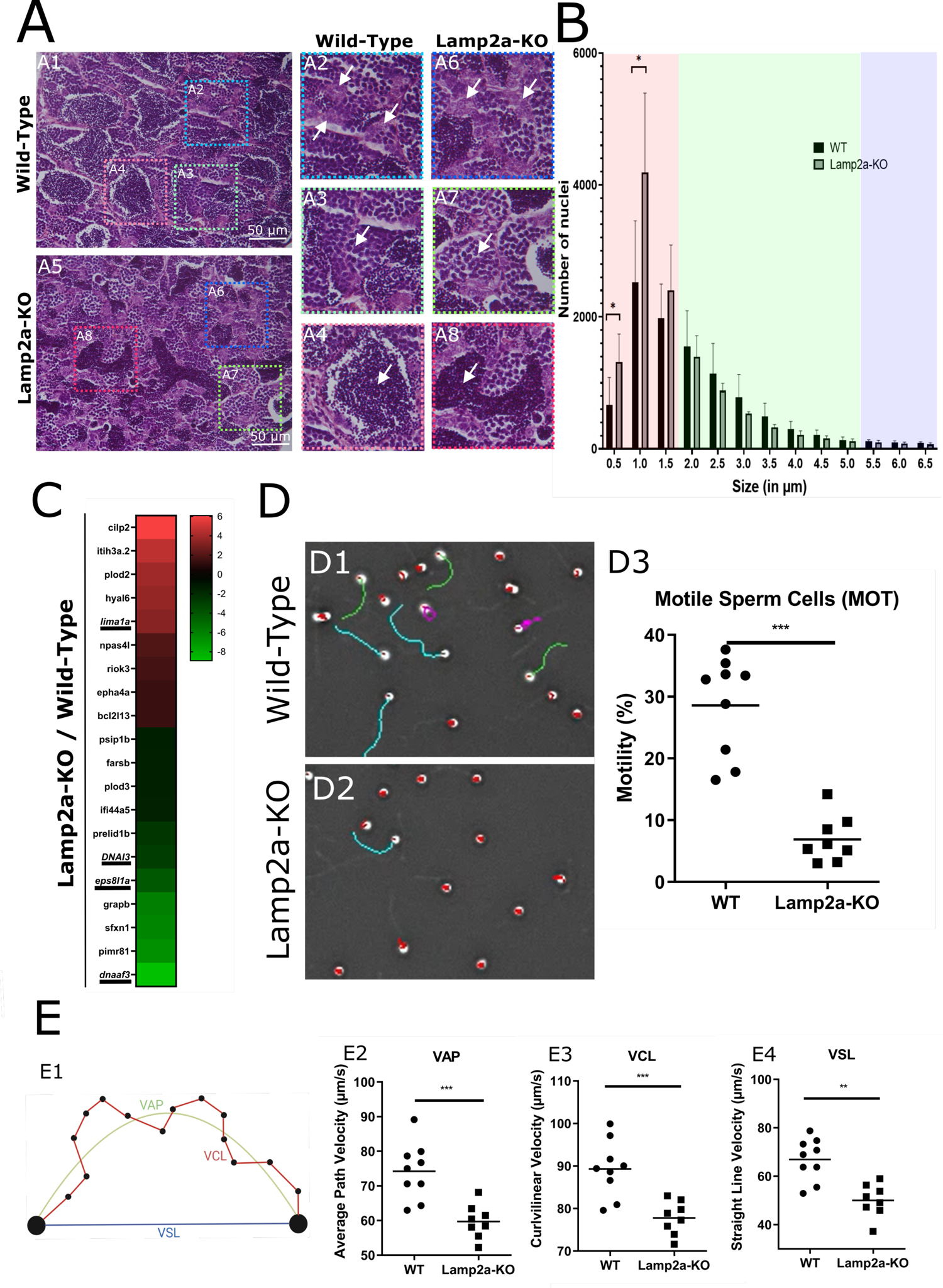
CMA controls the production of active spermatozoa. (A) Testes from WT (A1-A4) and Lamp2A-KO (A5-A8) zebrafish were stained with H&E. A2-A8 show boxed areas at higher magnification. (B) Quantification of the number of nuclei in each size category. For each size category the different lines (WT vs Lamp2a-KO) were compared with a t-test (p<0.05*). (C) Heatmap of RNA-seq data depicting differences in expression levels of selected genes between Lamp2a-KO and WT zebrafish. Four independent samples for each lines were used for RNA-seq experiment. (D) Representative images from videos of semen samples from WT (D1) and Lamp2a-KO (D2) zebrafish analyzed with CASA. Blue tracks = rapid-swimming sperm, green tracks = medium-swimming sperm, pink tracks = spermatozoa spinning on themselves, red tracks = static sperm. D3, Percentage of motile sperm cells from WT and Lamp2a-KO zebrafish. Each reported value corresponds to one fish (for a total of n=8-9 fish per line, corresponding to a final number of 3200-3600 spermatozoa analyzed per line). The horizontal lines represent the average for each condition. The different conditions were compared with a t-test (p<0.001***). (E) Analysis of sperm motility parameters in WT and Lamp2a-KO zebrafish. E1, Detailed kinematics measured for each sperm cells. Curvilinear velocity (VCL) is the distance over time of the actual path of the sperm head, whereas path velocity (VAP) is the distance along a 5 point mathematically smoothed path that the sperm travels over time. The third velocity measurement “straight line velocity” (VSL) is the straight line distance between the position of the sperm head in the first frame of analysis and the position of that sperm head in the last frame of analysis over the fixed time of analysis. E2-E3, Each reported value corresponds to one fish (for a total of n=8-9 fish per line, corresponding to a final number of 3200-3600 spermatozoa analyzed per line). The horizontal lines represent the average for each condition. The different conditions were compared with a t-test (p<0.001***).

To unveil how Lamp2a deficiency could underlie the profound phenotypic changes observed, we performed a quantitative proteomic analysis of sperms from either WT or Lamp2a-KO fish. Overall, we identified 2887 proteins, among which 6.03% (174 proteins) were significantly downregulated and 3.53% (102 proteins) were significantly upregulated in Lamp2a-KO sperm compared to WT (Figure 5A1 and Tables S2 and S3). Gene Ontology analysis revealed that the most downregulated biological processes are associated with negative regulation of peptidase activity and lipid transport (Figure 5A2). Upon closer examination, we observed that the downregulated proteins predominantly comprise a subset of proteins identified as acute phase proteins (APPs), which are often implicated in inflammatory responses^40,41^ and include proteinase inhibitors (e.g., SERPIN- and FETUIN-families member proteins), apolipoproteins, complement components, transferrin, and hemopexin (Figure 5B and Table S2). Given the pivotal role that APPs play in safeguarding sperm integrity^42^, the diminished expression of these proteins in spermatozoa from Lamp2a-KO, in comparison to the control group, might account for the observed compromised sperm quality in the CMA-deficient fish. Interestingly, the Gene Ontology analysis also highlighted that the most upregulated process relates to the mitochondrial respiratory chain (Figure 5A3). Indeed, several subunits of mitochondrial Complex I (NADH dehydrogenase [ubiquinone] 1 alpha subcomplex subunit 6 (NDUA6)), complex III (Cytochrome b-c1 complex subunit 8 (UQCRQ), Cytochrome b-c1 complex subunit Rieske (UQCRFS1)) and complex IV (cytochrome c oxidase subunit 4 (COX4), cytochrome c oxidase subunit 6A1 (COX6A1)), are found to be accumulated upon inactivation of Lamp2A (Figure 5B and Table S3). Notably, complement component 1 Q binding protein (C1QBP), which has previously shown to be critical in supporting translation of the mitochondrially encoded respiratory chain protein complexes^43^, was also over-represented in spermatozoas of Lamp2a-KO fish. These results *de facto* support the hypothesis that the low Lamp2a-KO sperm motility observed might be related to perturbations in the mitochondrial respiration chain process. To test this assumption, we next quantified the mitochondrial membrane potential (MtMP), a global indicator of mitochondrial’s function and integrity. Sperm MtMp was quantified using the membrane-permeant JC-1 dye and flow cytometry (Figures 5C and 5D1-5D2). Results indicated a significantly lower total MtMP in Lamp2a-KO spermatozoas compared to their WT counterparts (Figure 5E). Conversely a higher percentage of depolarized mitochondrias was evidenced in the former group compared to the latter (Figure 5F), supporting that the low Lamp2a-KO sperm motility observed may indeed be related to perturbations in mitochondrial respiration. Collectively, data gathered by quantitative proteomic analysis provide compelling evidence that the absence of CMA-activity indeed triggers substantial reconfigurations within the spermatozoa proteome. Furthermore, they suggest a pivotal role for CMA in governing mitochondrial metabolism within sperm cells. Notably, using the KFERQ finder V0.8 application^44^ to examine CMA-targeting motifs within the sequences of the 102 proteins up-regulated in Lamp2a-KO spermatozoa, we found that most (67%) mitochondria-related proteins display at least one KFERQ-like motif (Figure 5G1 and Table S4), underscoring the plausible direct engagement of CMA in orchestrating their turnover. Similarly, 80% of the other (non-mitochondrial) up-regulated sperm proteins exhibited at least one KFERQ-like motif (Figure 5G2 and Table S4), bringing them as potential CMA substrates candidates. Overall, these results emphasize a previously unknown and critical function for CMA as a gatekeeper of sperm cells proteostasis and quality.

**Figure 5.**
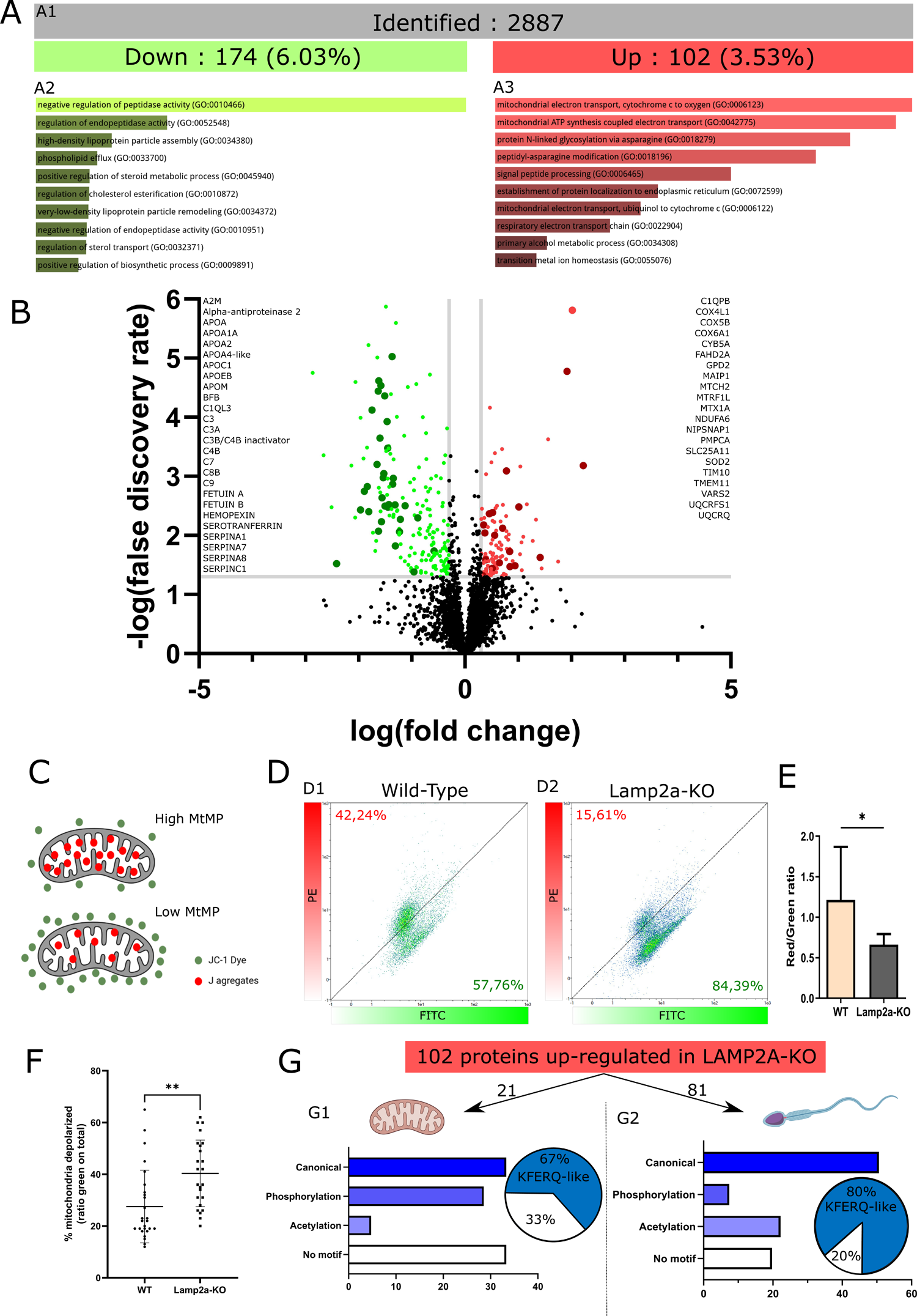
Deletion of Lamp2a in zebrafish leads to substantial remodeling of the sperm proteome and to defects in mitochondrial functions. (A and B) Quantitative proteomic analysis of sperm from WT and Lamp2a-KO fish. (A1) Number of identified proteins. (A2 and A3) Number and percentage of significantly down- and up-regulated hits in sperm from Lamp2a-KO *VS* WT zebrafish. (B) Volcano plot of the quantitative proteomics analysis of sperm from Lamp2a-KO *VS* WT zebrafish. Green and red dots indicate significantly down- and up-regulated proteins, respectively (p<0.05 and fold change >1.25). Protein list on the left: Subset of down-regulated proteins identified as acute phase proteins. Protein list on the right: Subset of up-regulated proteins involved in the mitochondrial respiratory chain. (C) Schematic illustration depicting JC-1 entry into the mitochondria and the generation of J aggregate. JC-1, a cationic carbocyanine dye (green) exhibits potential dependent accumulation in mitochondria where it starts forming J aggregates (red); upon depolarization, it remains as monomer showing green fluorescence. The change in ratio of red to green fluorescence is used as an indicator of mitochondrial condition. (D) Representative flow cytometry analysis of JC-1 staining in sperm cells from WT (D1) and Lamp2a-KO (D2) zebrafish. Horizontal (green) axis and Vertical (red) axis represent fluorescence intensities of JC-1 monomers and JC-1 aggregates, respectively. Increased intensity of monomers indicates decreased MtMP. (E) Ratio of JC-1 monomers and aggregates is presented as the Mean ± SD of 3 independent experiments. Ratios were significantly decreased in Lamp2a-KO sperm cells compared to their WT counterparts. *p<0.05, tested by Student t-test. (F) Percentage of depolarized mitochondria calculated by the ratio of green fluorescence to the total fluorescence and presented as the Mean ± SD of 3 independent experiments. **p<0.01, tested by Student t-test. (G) Proportion of up-regulated proteins related to mitochondria (G1) or to other (non-mitochondrial) functions (G2) harboring or not distinct classes of KFERQ-like motifs. The pie charts show the proportion of all proteins with at least one KFERQ-like motif (blue) *VS* those with no motif (white).

## DISCUSSION

Recently, using two distinct fish species: (i) the medaka fish model (*Oryzias latipes*, as reported by Lescat et al. in 2020^23^), and subsequently (ii) the rainbow trout (*Oncorhynchus mykiss*, as documented by Vélez et al. in 2023^26^), we extended the spread of a conserved functional CMA activity to teleosts in complement to mammals and birds. These discoveries represented a significant milestone in our understanding of the evolutionary history of CMA, shedding light on its presence and functionality from the onset of vertebrates. Dysregulations or malfunctions of CMA have been linked to the causative development of various human pathologies, including neurodegenerative diseases, cancers, and immune disorders^1^. Understanding the role of CMA in fish, which share evolutionary and physiological similarities with humans, offers a promising platform for elucidating the molecular mechanisms underlying these pathologies.

Building on our first reports on the evolutionary radiation of the CMA function in vertebrates, we have then stepwise demonstrated that CMA is not limited to medaka and trout, but is also operative in a widely studied model organism: the zebrafish (*Danio rerio*). This demonstration does not only broadens our understanding of this essential cellular process in another fish species, but furthermore capitalizes on the unique advantages this model organism offers: low cost feeding and maintenance, large numbers of offspring, rapid development *ex utero*, amenability to genetic and chemical screens, optical transparency of embryos and larvae which facilitates noninvasive visualization of organs and biological processes *in vivo*^45^.

In the present study, a significant step forward has been accomplished after we were able to develop different zebrafish transgenic lines (KFERQ-PS-Dendra2 and KFERQ-PA-mCherry) tailored for the precise monitoring of CMA activity within tissues at the cellular resolution. Through image-based analyses, we were able to demonstrate the presence of fasting-inducible CMA activity in specific cell types, such as renal tubules - as previously reported in the KFERQ-Dendra mouse model^19^ - or unexpectedly, in the Sertoli cells of testes. These results therefore attest to the virtually unrestrained potential of these zebrafish CMA reporter lines as a complementary model to the KFERQ-Dendra mouse for studying this function in organs with complex cellularity.

Thus, the usage of the zebrafish model together/combined with the tools developed for monitoring and assessing the CMA function allowed us to uniquely highlight the so far underappreciated involvement of CMA during spermatogenesis and its decisive role for sperm motility. As previously mentioned, our image-based analyses of the CMA-reporter zebrafish line unveiled the presence of discrete yet highly specific CMA activity within a particular cell type in male gonads, referred to as Sertoli cells. Given the pivotal role of these cells in supporting and nurturing germ cell development in the adult gonads^46^, we conducted an evaluation/measurement of spermatozoa ‘quality’ in Lamp2a-KO zebrafish in comparison to the WT. Results revealed a significant reduction in sperm motility in Lamp2a-KO fish when compared to their WT counterparts. Proteomic analyses then suggested that defects in the mitochondrial respiratory chain may underlie, at least in part, this effect. Mitochondria play a pivotal role in germ cell development and any defects in mitochondrial quality can have serious consequences on male fertility^47^. Into that direction, several studies established clear correlations between sperm motility and both MtMP and the mitochondrial respiratory complex activities^48^. Since mitochondria are the major source of pro-oxidative agents, it is suggested that dysfunction of this organelle would have a fundamental role in the oxidative imbalance affecting sperm function^49^. Wang et al. thus identified low MtMP and high reactive oxygen species (ROS) production in spermatozoa from infertile patients, probably as a consequence of such mitochondrial injury^50^. Mitochondria are also critical for sperm metabolism and energy production^51^, and defects in mitochondria have been shown to affect energy furniture for sperm motility^52^. Collectively, our data designate mitochondria as the obvious key organelle for handling sperm motility. In this context, the decreased sperm motility observed in Lamp2a KO fish could be related, at least partly, to defects of mitochondrial activity. Recent research has already highlighted the role of CMA in controlling the levels of several proteins involved in mitochondrial functions^53–55^. However, to the best of our knowledge, our findings represent the first evidence that this effect could affect male reproduction. Future research will be essential to further elucidate this relationship between CMA, mitochondria and sperm motility.

In summary, this article approaches the study of CMA from an entirely new and innovative angle, spanning from the initial discovery of CMA activity in zebrafish to the development of a specialized transgenic line - that allows determining and tracking the spatiotemporal dynamic changes of CMA activity *in vivo* - and culminating in the revelation of CMA’s pivotal role in reproductive processes. Overall, our work not only expands the frontiers of knowledge surrounding CMA but also underscores its relevance in a broader biological context, with potential implications for both fundamental research and reproductive health studies.

## MATERIALS AND METHODS

### Zebrafish breeding and treatments

Adult zebrafish male and female were used. Fish were grown at 27°C under a 12 h photoperiod time (12 h in light, 12 h in dark). They were fed 3 times a day with commercial breeding. During treatments, fish were starved for 48 h. For organs sampling, zebrafish were euthanized in a lethal dose of Tricaïne (MS-222). All experimental procedures were conducted in strict accordance with the legal frameworks of France and the European Union. They respect the directive 2010/63/EU relating to the protection of animals used for scientific purposes as well as the decree No 2013-118, February 1, 2013, of the French legislation governing the ethical treatment of animals.

### Construction of CMA reporter plasmids and design of morpholino antisense oligonucleotide

The PA-mCherry1-N1 vector was a gift from Michael Davidson (Addgene, 54507; http://n2t.net/addgene:54507; RRID: Addgene_54507; deposited by Michael Davidson), and from this we generated a KFERQ-PA-mCherry1 construction composed by the N-terminal 21 amino acids of bovine RNASE1/RNase A containing its KFERQ CMA-targeting motif, fused to the photoactivable-mCherry1 (PA-mCherry1) protein, as originally reported by Koga et al (Koga et al., 2011). The functionality of our construct has recently been validated in fish cells (Lescat et al., 2020; Vélez et al., 2023). The pKFERQ–Dendra2 plasmid was constructed by inserting the oligonucleotide coding for the N-terminal sequence of the bovine ribonuclease A containing the KFERQ-CMA targeting motif (MKETAAAKFERQHMDSSTSAA) into the p3167 vector (gifted from Pr Larazo Centanin in Centre of Organismal Studies in Heidelberg, Germany). The LAMP1-GFP construct was a gift from Benjamin Dehay (Univ. Bordeaux, INSERM, France).

Anti-ATG7 Morpholino oligonucleotide (Mo) (AGCTTCAGACTGGATTCCGCCATCG) was previously descripted in (Lee and al., 2014) and designed with Gene Tools, to directly target Atg7 protein. Embryos were injected at the 1-cell stage with 1 mM. Mo action were confirmed by Atg5/Atg12 formation in cells by Western blotting.

### CMA activity assay in zebrafish primary embryonic cells

To track and quantify CMA activity, embryos were micro-injected with a KFERQ-Dendra2 RNA, KFERQ-PA-mCherry or Lamp1-GFP during one cell stage. After 8 h of development, at 50% epiboly, cells were isolated and plated into culture in DMEM (Dulbecco’s Modified Eagle Medium), with 10% FBS (Foetal Bovine Serum) and 5% antibiotics (Streptomycine). During the treatments, cells were cultured at 25°C and were incubated in a serum-free DMEM (Serum-) or in combination with 25 µM of H_2_O_2_. Then, cells were photoconverted by a 405 nm laser for 10 minutes at 2.80 A, and cells were then maintained in medium culture for 16 h. To check for lysosomal translocation, protease inhibitors were used, with 50 nM Leupeptin and 50 nM NH_4_Cl.

### Establishment of Zebrafish KO and transgenic lines

Zebrafish deleted for Lamp2a were generated using the CRISPR/Cas9 method. Guide RNAs (sgRNAs) targeting two sites located before and after Lamp2a exon were designed using ZiFiT software (http://zifit.partners.org/ZiFiT/Disclaimer.aspx). DR274 vector (Addgene #42250) containing the guide RNA sequence was first linearized with Bsa1, electrophoresed in a 2% agarose gel and purified. PCR amplifications were then performed using linearized DR274 as a template and two primers for each sgRNA. Forward primers containing sgRNAs target sequences (in red and bolded in Figure S1C). Forward primer: 5’-GAAATTAATACGACTCACTATAgtgtaaatgtcactaaagtcGTTTTAGAGCTAGAAATAGCA AG-3’. Reverse primer: 5’-GAAATTAATACGACTCACTATAgttctgttgtgatctaacatGTTTTAGAGCTAGAAATAGCAAG-3’. The sgRNAs were co-injected with Cas9 protein. Synthesized RNAs were then injected into 1-cell stage zebrafish embryos at the following concentrations: 25 ng/mL for each sgRNA and 100 ng/mL for the Cas9 protein. Genotyping was performed on gDNA from caudal fin-clips of adult fishes. CRISPR-positive fish were screened for mutations using a set of PCR primers (Forward primer: 5’-CGTCAATTTACTGCTACTCTAGCC-3’; Reverse primer: 5’-TTATACAGACTGGTAACCAGTACG-3’) flanking the sgRNAs target sites leading to a 737 bp deletion on the exon 6A of the *Lamp2* gene (Figures S1B and S1C).

KFERQ-PS-Dendra2 and KFERQ-PA-mCherry zebrafish reporter lines were generated by injecting the previously described plasmids into 1-cell zebrafish embryos. These plasmids contain a crystalline Beta cassette, which was used to select fish that had integrated the plasmid by fluorescence in the eye. Once adult, these fish were crossed with each other to obtain global reporter fish due to the ubiquitin promoter. The Tg(*gsdf*:eGFP);KFERQ-PA-mCherry zebrafish line was generated by crossing Tg(*gsdf*:eGFP) transgenic zebrafish with the newly generated KFERQ-PA-mCherry zebrafish reporter line.

### Microscopy imaging

Images of cells were taken in the MRic Photonics platform (Biosit Rennes) with a DeltaVision Elite high-resolution microscope equipped with a 60X oil objective lens (GE Healthcare). Images were spliced into different channels and we applied a filter (plugin) with Image J (https://fiji.sc/ RRID:SCR_002285) to directly quantify the puncta from the reporter. Images from tissues were taken using a LEICA confocal microscope SP5 with a 40X oil objective lens. Images were processed using LAS-AF software (Leica Microsystems).

### Lysosomal content

Acidic vesicles were controlled between the Wild-Type and Lamp2a-KO cells using either Lamp1-GFP by microinjection in a one cell stage zebrafish embryo, or by incubating cells and tissues in Lysotracker Deep Red DND-26 (Invitrogen) for 30 minutes at 75 nM, and were then observed by microscopy and quantify by Image J.

### Sperm motility

Male Zebrafish were isolated the day before sampling. The day after, they were anesthetized and sperm sampling were done directly by aspiration. Sperm were then diluted in HBSS (Hanks’ balanced salt solution) at 300 µM and conserved in ice to reduce their motility. They were displayed in the CASA (Computer Assisted Sperm Analysis) and motilities were recorded using the module IVOS II (IMV technology) and the camera ISAS 782M camera recorder. Range size particles were defined between 2 and 20 mm in the CASA settings.

The parameters considered in this study were: total motility (MOT, %); the percentage of fast (average path velocity [VAP] >100 mm/s), medium (VAP ¼ 50–100 mm/s), and slow (VAP ¼ 10–50 mm/s) spermatozoa; curvilinear velocity (VCL, in mm/s), defined as the time/average velocity of a sperm head along its actual curvilinear trajectory; straight line velocity (VSL, mm/s), defined as the time/average velocity of a sperm head along the straight line between its first detected position and its last position; VAP (mm/s), defined as the time/average of sperm head along its spatial average trajectory; straightness (STR, %), defined as the linearity of the spatial average path, VSL/VAP^56^. At least 400 spermatozoids were counted in each sample.

### Mitochondrial Assay

Male Zebrafish were isolated the day before sampling. The day after, they were anesthetized and sperm sampling was done directly by aspiration. Sperm were then diluted in a HBSS solution at 300 µM and conserved in ice. Then, sperm were incubated with 50 nM of JC-1 labelling (Invitrogen) for 15 minutes at 20°C. Control was performed by adding the same amount of DMSO in sperm. They were observed in flux cytometry and each sample was measured in triplicate with 3 different experiments.

### Histological preparations

Testis sampling were fixed in Bouin at 4°C for overnight under shaking. After several dehydration bath of croissant ethanol concentrations (25%, 50%, 70%, 96%), tissues were flushed in PBS, covered in paraffin and then sectioned in 5 nm by microtome. The sections were mounted in glasses and stained with hematoxylin and eosin. The various cell stages were counted using Fiji and the Stardist plugin. Score threshold was set to 0.5 and the overlap threshold to 0.4.

### Quantitative RT-PCR analysis

For RNA extraction, tissues from 6 different zebrafish were pooled in each condition and we used TriZol protocol to extract total RNA (Invitrogen, Carlsbad, California, USA). The protocol conditions for sample preparation and quantification have been previously detailed (Seiliez and al, 2016). The primer used for RT-PCR were listed in table S5. The relative quantification of target genes expression was calculated using the deltaCT method and 18S was used as housekeeping gene. Samples were analysed in triplicates.

### Bulk RNA barcoding and sequencing (BRB-seq) analysis

Testes were collected from wild type and Lamp2a-KO adult males, under fed and fasting conditions. We used the TriZol protocol to extract RNA. After Nanodrop quantification, the concentration of each sample was adjusted to 50 ng. Samples were then sent to Alithea Genomics S.A. First, the RNAs were barcoded and then retrotranscribed. Once synthesized, the second strand is synthesized. These cDNA strands were then segmented and labelled. From these DNA fragments, the fragments were amplified and 2 libraries were created. The first contains only barcode sequences and unique molecular identifiers (UMIs). The second contains all the RNA fragment sequences produced by Illumina. From here, the sequencing data are analysed, the UMIs are counted and a gene-counting matrix is set up. Next, a demultiplexing step is required to match the codes in the first library with the sequences in the second, and also with the reference zebrafish genome, and was done by R script. Significance level was set at p < 0.05, and ratios (Fold change, FC) were considered relevant if higher than +/- 1.

### Protein extraction, western blotting and quantitative proteomics

Proteins were collected using RIPA buffer (ThermoFisher Scientific, 89901) supplemented with protease and phosphatases inhibitor cocktail (ThermoFisher Scientific, 78422). Protein concentration was determined using the Qubit protein assay kit (ThermoFisher Scientific, Q33211).

For Western blot analyses, the samples were then electrophoresed by sodium dodecyl sulfate polyacrylamide gel electrophoresis (SDS-PAGE). Afterwards, proteins were transferred to polyvinylidene fluoride (PVDF) membranes (Merk-Millipore, IPFL00010), blocked with SuperBlock Blocking Buffer in PBS (Thermo Scientific, 37515), and immunoblotted following the manufacturer’s protocol using an antibody against Atg5 (Novus, NB110-53818). No-Stain™ total protein labelling was used as a loading control. The different immunoreactive bands were developed using the SuperSignal West Pico Plus Chemiluminescent Substrate (ThermoFisher Scientific, 34578). Signal acquisition was performed using Smart Exposure in an iBright FL1500 Imaging System (ThermoFisher Scientific, Illkirch Cedex, France) to assure signal linearity, and then images were quantified with the iBright Analysis Software (ThermoFisher Scientific, RRID:SCR_017632).

For quantitative proteomic analysis, 50 µg of protein extract from a pool of 5 individual fish with 5 replicates per condition were extracted and shipped to the ProteoToul Services - Proteomics Facility of Toulouse (https://proteotoul.ipbs.fr/proteotoul-services/) for posterior analysis. Briefly, 50 µg of dried protein extracts from each sample were solubilized with 25 µl of 5% SDS. Proteins were submitted to reduction and alkylation of cysteine residues by addition of TCEP and chloroacetamide to a final concentration respectively of 10 mM and 40 mM. Protein samples were then processed for trypsin digestion on S-trap Micro devices (Protifi) according to manufacturer’s protocol, with the following modifications: precipitation was performed using 216 µl S-Trap buffer, 4 µg trypsin was added per sample for digestion in 25 µl ammonium bicarbonate 50 mM. After that, tryptic peptides were resuspended in 35 µl of 2% acetonitrile and 0.05% trifluoroacetic acid and analyzed by nano-liquid chromatography (LC) coupled to tandem mass spectrometry (MS), using an UltiMate 3000 system (NCS-3500RS Nano/Cap System; ThermoFisher Scientific) coupled to an Orbitrap Exploris 480 mass spectrometer equipped with a FAIMS Pro device (ThermoFisher Scientific). 1 µg of each sample was injected into the analytical C18 column (75 µm inner diameter × 50 cm, Acclaim PepMap 2 µm C18 ThermoFisher Scientific) equilibrated in 97.5% solvent A (5% acetonitrile, 0.2% formic acid) and 2.5% solvent B (80% acetonitrile, 0.2% formic acid). Peptides were eluted using a 2.5% - 40% gradient of solvent B over 62 min at a flow rate of 300 nL/min. The mass spectrometer was operated in data-dependent acquisition mode with the Xcalibur software. MS survey scans were acquired with a resolution of 60,000 and a normalized AGC target of 300%. Two compensation voltages were applied (−45 v/-60 v). For 0.8 s most intense ions were selected for fragmentation by high-energy collision-induced dissociation, and the resulting fragments were analyzed at a resolution of 30,000, using a normalized AGC target of 100%. Dynamic exclusion was used within 45 s to prevent repetitive selection of the same peptide. Thereafter, raw mass spectrometry (MS) files were processed with the Mascot software (version 2.7.0) for database search and Proline55 for label-free quantitative analysis (version 2.1.2). Data were searched against zebrafish entries of Uniprot protein database. Carbamidomethylation of cysteines was set as a fixed modification, whereas oxidation of methionine was set as variable modifications. Specificity of trypsin/P digestion was set for cleavage after K or R, and two missed trypsin cleavage sites were allowed. The mass tolerance was set to 10 ppm for the precursor and to 20 mmu in tandem MS mode. Minimum peptide length was set to 7 amino acids, and identification results were further validated in Proline by the target decoy approach using a reverse database at both a PSM and protein false-discovery rate of 1%. For label-free relative quantification of the proteins across biological replicates and conditions, cross-assignment of peptide ions peaks was enabled inside group with a match time window of 1 min, after alignment of the runs with a tolerance of +/- 600 s. Median Ratio Fitting computes a matrix of abundance ratios calculated between any two runs from ion abundances for each protein. For each pair-wise ratio, the median of the ion ratios is then calculated and used to represent the protein ratio between these two runs. A least-squares regression is then performed to approximate the relative abundance of the protein in each run in the dataset. This abundance is finally rescaled to the sum of the ion abundances across runs. A Student T-test (two-tailed t-test, equal variances) was then performed on log2 transformed values to analyze differences in protein abundance in all biologic group comparisons. Significance level was set at p < 0.05, and ratios (Fold change, FC) were considered relevant if higher than +/- 1.25.

The list of proteins identified in the present study is provided in Tables S2 abd S3 along with the FC (log2 transformed) and p-value (log10 transformed) results of the down- and up-regulated proteins in Lamp2a-KO vs WT fish. Gene Ontology enrichment maps were then generated using Enrichr^57,58^. Finally, the presence of KFERQ motifs within the protein sequences was assessed using the KFERQ finder app V0.8^44^, and data is presented in Table S4.

### Statistical analysis

All data are reported as means ± SD or means of percentages ± SD from a minimum of 3 independent experiments. Normality of data residuals were verified before performing parametric or non-parametric statistical tests. Except in Figure S4A for which Exact binomial tests were used, in all other cases, for comparison with more than 2 groups, one-way ANOVA analysis were performed followed by Tukey’s multiple comparison (same sample size) or Bonferroni’s multiple comparison (different sample size) post-hoc tests. For comparison between two groups, we used a parametric two-tailed unpaired Student’s T-test, or a non-parametric Mann Whitney test. All statistical analyses were performed using GraphPad Prism version 8.0.1 for Windows (GraphPad Software, Inc., www.graphpad.com) and a p-value < 0.05 was set as a level of significance.

## END NOTES

**Supplementary information** is available in the online version of the paper

## Acknowledgements

We thank S. Dutertre and X. Pinson from the Microscopy Rennes Imaging Center (MRic, BIOSIT, Biogenouest) for assistance. MRic is member of the national infrastructure France-BioImaging supported by the French National Research Agency (ANR-10-INBS-04). We also thank C. Labbé (from INRAE UR1037 Laboratory of Fish Physiology and Genomics) for their technical assistance. We acknowledge Alexandre Stella from the ProteoToul Services - Proteomics Facility of Toulouse for the proteomics analysis. We are grateful to J.J. Lareyre for kindly providing us with the Tg(*gsdf*:eGFP) zebrafish. We also thanks L. Peron for manufacturing the light emitting device. This research was supported by the Communauté d’Agglomération Pays Basque (CAPB).

## Author Contributions

A.H. and I.S. got the funding. M.G., A.H. and I.S. conceived and planned the study as well as all the experiments. M.G., E.J.V., S.S., V.V, K.D., and A.D. performed the experiments. M.G., A.H. and I.S. analyzed the data. M.G. and I.S. wrote the original draft of the manuscript. I.S., A.H., F.B., E.J.V., and S.S. reviewed and edited the final version of the manuscript. I.S. and A.H. supervised the project and had primary responsibility for final content.

## Data availability

The authors confirm that the data supporting the findings of this study are available within the article [and/or] its supplementary materials.

---

**Figure S1.** Generation of Lamp2a-Ko zebrafish using Crispr-cas9 method. (A) Schematic representation of the strategy used to generate the specific lamp2a knockout zebrafish. (B) Genotyping of *lamp2a* allele was performed by PCR of fin from fish generated by Crispr-cas9 method. Mutant fish (-/-) displayed a deletion of the 737bp (low band) compared to wild type (+/+) fish (high band). Heterozygous fish (+/-) are shown as an additional control. (C) Sequence alignment of the gene *lamp2* between wild type and mutant zebrafish showing the deletion in KO fish of the 737bp region in exon 6 (in green) that encodes for the specific cytosolic and transmembrane domains of the Lamp2a protein. (D) *lamp2a*, b and c transcripts levels obtained by RT-qPCR in whole fish, testis, gut and kidney of wild type and Lamp2a-KO zebrafish. Data are means ± SD. The data were normalized by the values of *18S*. A Student t-test was performed (p<0.05). ns, not significant.

**Figure S2.** KFERQ-Dendra2 reporter puncta generation does not require macroautophagy. (A and B) Atg12–Atg5 autophagy complex formation after anti-atg7 morpholino (Mo) injection. Embryos were injected at the 1-cell stage with control or anti-atg7 Mos, and lysates were prepared for immunoblot analysis at 16 hpf. (A) Immunoblot with anti-Atg5 antibody. No-Stain™ total protein labeling is shown as a loading control. (B) Densitometric analysis. Data are expressed relative to control (WT) zebrafish (n=1 pool of 50-60 embryos/condition). (C) Representative images of zebrafish primary embryonic cells micro-injected with both mRNAs coding for the KFERQ-Dendra2 reporter and anti-atg7 Mo. After photoactivation, cells were maintained in the presence (Serum +) or absence of serum (Serum-) for 16 h. Nuclei were stained with Hoescht 33342. (D) Quantification of KFERQ-Dendra2 reporter red puncta per cell. All values correspond to individual images (n=50 cells/condition). The lines represent the average for each condition. The two conditions were compared with a t-test (p<0.001***).

**Figure S3.** RT-qPCR for Dendra in the indicated tissues relative to the average of *18S* in wild type (WT) and Lamp2a-Ko fed and starved zebrafish. Data are means ± SD. A Student t-test was performed (p<0.05). ns, not significant.

**Figure S4.** Expression of the gene *LAMP2* (from single nucleus RNA profiling) along spermatogenesis for human, mouse, opossum, and chicken. Exact binomial tests were performed for statistical comparisons (****P< 0.0001). OS, Other somatic; ST, Sertoli; SG, Spermatogonia; SC, Spermatocytes; rSD, Round spermatids; eSD, Elongated spermatids.

**Video S1.** Video showing all the z-stack images acquired from representative tubules of KFERQ-Dendra2 WT zebrafish starved for 48 h.

**Video S2.** Video showing all the z-stack images acquired from representative tubules of KFERQ-Dendra2 -Lamp2a-KO zebrafish starved for 48 h.

**Video S3.** Video of sperm from WT zebrafish analyzed with CASA. Blue tracks = rapid-swimming sperm, green tracks = medium-swimming sperm, pink tracks = spermatozoa spinning on themselves, red tracks = static sperm.

**Video S4.** Video of sperm from Lamp2a-KO zebrafish analyzed with CASA. Blue tracks = rapid-swimming sperm, green tracks = medium-swimming sperm, pink tracks = spermatozoa spinning on themselves, red tracks = static sperm.

