## Supplementary figures and images for "The zebrafish as a new model for studying chaperone-mediated autophagy unveils its role in spermatogenesis"

### Supplemental Figures

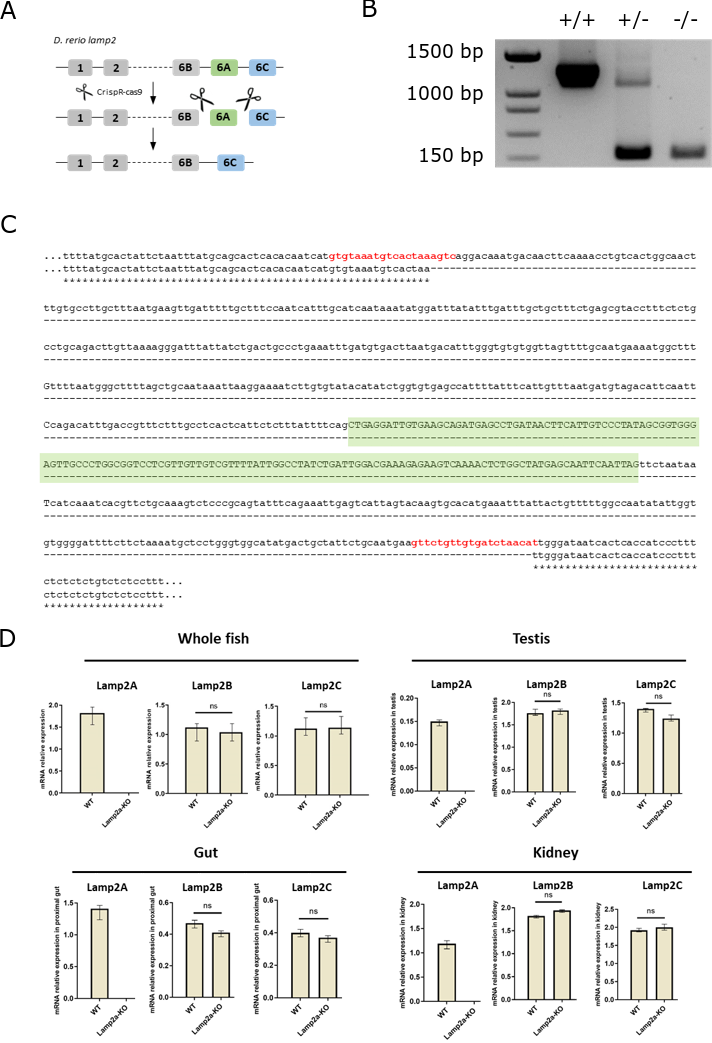

**Figure S1**

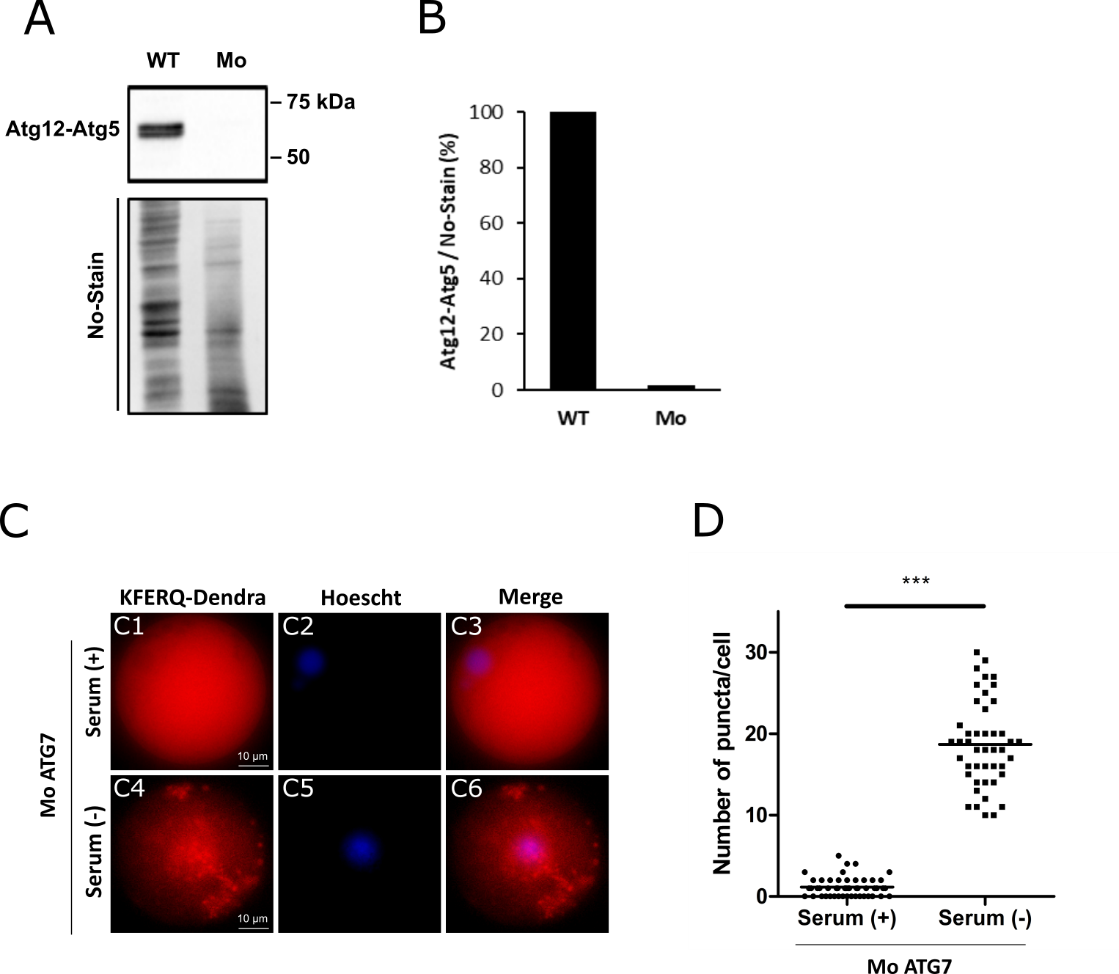

**Figure S2**

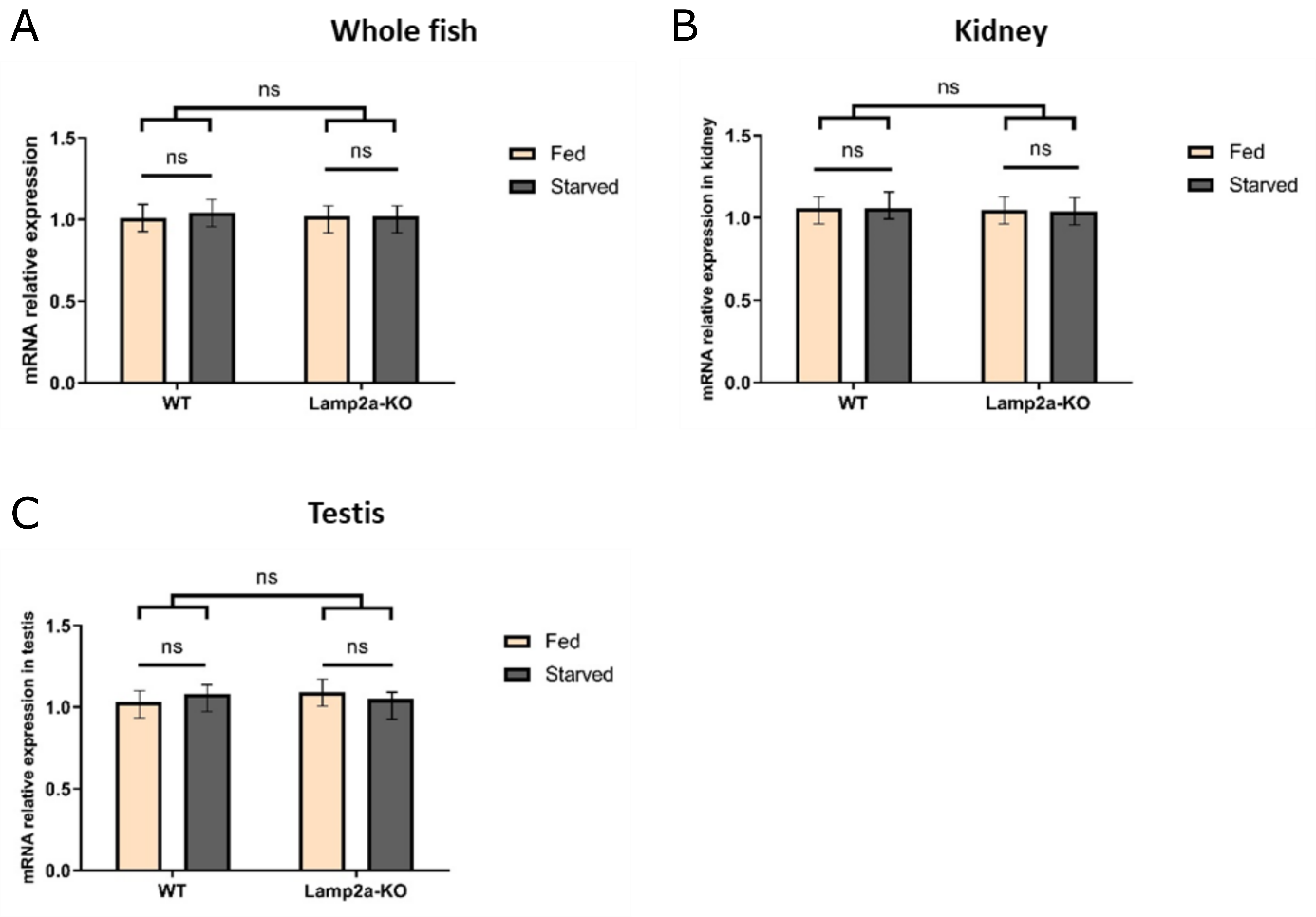

**Figure S3**

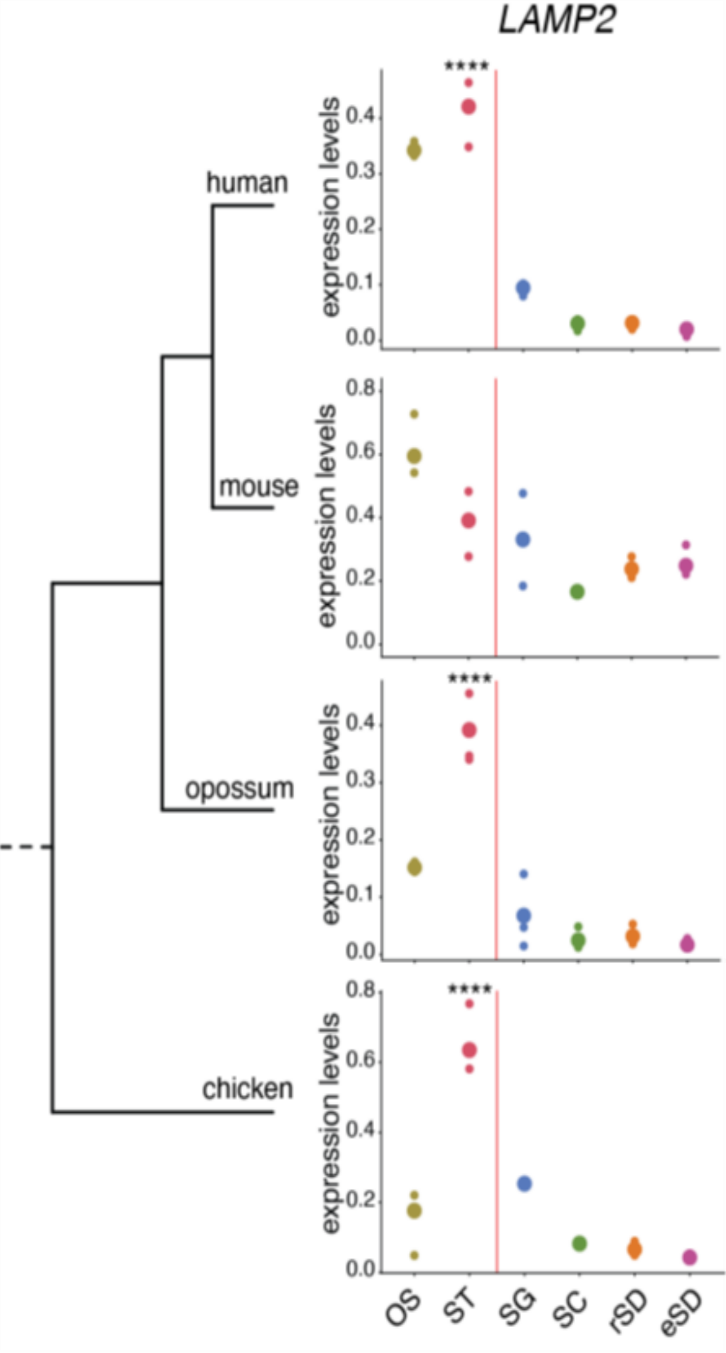

**Figure S4**
